# Gelatin coating enhances flow diverting stent endothelial cell coverage, parent vessel healing and aneurysm occlusion

**DOI:** 10.64898/2026.07.30.741814

**Authors:** John W. Thompson, Jennifer S. Suon, Naser Hamad, Gina Corsaletti, Rianna Haniff, Sai Sanikommu, Maxon V. Knott, Pedro Bartkevitch Rodrigues, Helena Hernandez-Cuervo, Ahmed Abdelsalam, Tiffany A. Eatz, Jayro Toledo, Evan M. Luther, Robert M. Starke

## Abstract

**Background:** Flow diversion stent treatment of cerebral aneurysms has demonstrated high rates of aneurysm occlusion and long-term durability. However, complete endothelization of the flow diverting stent is required for parent vessel healing which closes the aneurysm and metal stent from the circulation. Therefore, stent coatings which enhance endothelial migration and attachment may increase the rate of aneurysm occlusion and reduce complications associated with flow diversion treatment. Here we investigate the use of gelatin as a stent surface coating to enhance the rate of stent endothelialization and coverage and increase aneurysm healing.

**Methods:** Nitinol- Neuroform stents (Stryker, Kalamazoo, MI) and cobalt-chromium-Pipeline Flex flow diverting stents (Medtronic, Minneapolis, MN) were used for this study. The stents were coated with gelatin and endothelial cell attachment, proliferation and stent coverage were determined *in vitro* and compared to uncoated stent controls. A rabbit elastase-aneurysm model was used to determine the effects of endothelial cell seeded-gelatin coated flow diverting stents on aneurysm obliteration and parent vessel healing.

**Results:** *In vitro*, gelatin coating of nitinol stents did not significantly alter endothelial cell attachment, proliferation, or stent coverage. However, gelatin coating of cobalt-chromium stents significantly increased endothelial cell attachment, proliferation and migration. In fact, gelatin coating significantly (p< 0.001) increased the rate of complete stent endothelization by 33% compared to unmodified controls. *In vivo*, treating aneurysm with endothelial cell seeded-gelatin coated stents resulted in aneurysm occlusion in 8 of 8 (100%) rabbit aneurysms at 90 days compared to only 4 of 7 (57%) in unmodified controls (p< 0.001). Histologically, there were trends in increased neoarterial wall thickness across the aneurysm neck and neointimal formation in the parent artery. Angiographic assessment demonstrated strong parent and side branch patency.

**Conclusions:** Gelatin coating enhances EC attachment and stent coverage which is dependent upon the type of stent. Endothelial cell seeded-gelatin coated-flow diverting stents allowed 100% aneurysm obliteration and neoarterial formation without affecting side branch patency or parent artery perfusion. Gelatin coating therefore represents a valuable strategy to enhance stent cellularization and aneurysm occlusion rates.

## BACKGROUND

Flow diversion has revolutionized the endovascular treatment of intracranial aneurysms, particularly wide based complex aneurysms that are challenging for endovascular or microsurgical techniques^1,2^. Multiple generations of flow diverters have demonstrated high rates of progressive aneurysm occlusion and low recurrence^3,4^. However, despite dual antiplatelet therapy, thromboembolic complications remain a significant concern due to the high metal coverage of flow diverters^1,5^. Additionally, the need to inhibit platelet function increases the risk of hemorrhagic complications^6,7^. Furthermore, aneurysms with continued filling despite flow diversion remain at risk of rupture and are particularly challenging to treat.

Flow diversion stent surface modifications have emerged to mitigate thrombogenicity while preserving or enhancing hemodynamic diversion and endothelial healing^8,9 10^. The majority of these surface modifications reduce thrombogenicity through antithrombotic drug and inhibitor coatings or by coating stents with hydrophilic biological mimics that shield the flow diverter from thrombogenic protein components^8,9,11,12^. There is limited evidence if these stent surface coatings improve aneurysm occlusion and limit thromboembolic complications.

Another but less investigated approach is the rapid endothelization of the stent thereby removing the device from the circulation. For example, coating coronary artery stents with anti-CD34 antibodies increase bind of circulating CD34+ endothelial progenitor cells and increased stent endothelialization^16–18^. Likewise, coating flow diverter stents with CD31 a transmembrane glycoprotein expressed on endothelial cells, platelets and lymphocytes, has been shown to decrease platelet and lymphocyte activation and improve endothelialization and aneurysm occlusion^19–21^.

Gelatin, a natural biopolymer derived from collagen hydrolysis, is a promising coating material for medical devices, particularly vascular and endovascular implants^22,23^. Gelatin is FDA approved and has excellent biocompatibility, biodegradability, low immunogenicity, and inherent capacity to mimic the extracellular matrix, thereby facilitating cell adhesion, proliferation, and tissue integration^24,25^. Gelatin-based modifications can be layered with anticoagulants like heparin or polydopamine, thereby improving antithrombotic properties and supporting neointimal healing in artificial vascular grafts and stent scaffolds^26,27^. These attributes position gelatin as an attractive, stent coating material for the rapid endothelialization of flow diverter stents. Therefore, in this study we investigated if gelatin coating would increase endothelial cell attachment and coverage on stents composed of nitinol or cobalt-chromium-nickel metals and if endothelial cell seeded-gelatin coated flow diverting stents enhance aneurysm occlusion and parent vessel healing.

## METHODS

All procedures were performed in accordance with institutional guidelines for animal research and approved by the University of Miami Institutional Animal Care and Use Committee (assurance number: A-3224-01).

### Materials

RPMI-1640 media, Fetal Bovine Serum (FBS), MEM amino acids, sodium pyruvate, and penicillin/streptomycin were purchased from Gibco/Life Technologies (Grand Island, NY). NuSerum, Hoechst 33342, Calcein-AM and propidium iodide were purchased from ThermoFisher Scientific (Waltham, MA). All other reagents were purchased from Sigma-Aldrich (St. Louis, MO) unless otherwise noted.

### Gelatin and Fibronectin Stent Coating

Standard Neuroform Atlas nitinol stents (Stryker, Kalamazoo, MI) and Pipeline Flex flow-diverting stents (Medtronic; Minneapolis, MN) were used for this study. Stents were coated in either 1% bovine gelatin dissolved in PBS or 1.5% Fibronectin diluted in PBS. Fresh coating solutions were made prior to use. Stents were coated by complete immersion in the gelatin or fibronectin solutions with gentle mixing overnight at 37°C. Control unmodified stents were placed in PBS alone. The devices were washed with PBS to remove excess gelatin or fibronectin and then used for experiments.

### Cell Culture

Human brain microvascular endothelial cells (ScienCell Research Lab; Carlsbad, CA) and GFP expressing human aortic artery ECs (Accegen, Fairfield, NJ) were cultured in RPMI-1640 media supplemented with 10% FBS, 10% NuSerum, 1% MEM amino acids, 1% sodium pyruvate, and penicillin/streptomycin. For experimentation purposes, the cells were plated in 100mm dishes and grown to 80% confluency. The cells were then trypsinized with trypsin-EDTA, centrifuged and the pellet resuspended in complete cell culture media at 5×10^5^ cells/ml.

#### Coating Stents with Cells

Coated and uncoated stents were placed in an open 1.5ml Eppendorf tube containing 1 ml of endothelial cell suspension. The tubes were transferred to a cell culture dish to maintain sterility and placed at 37°C with 5% CO2 for a total of 3 hours. The tubes were gently mixed every hour to resuspend the cells. The stents were then placed into a 100 mm culture dish containing PBS and unattached cells and cell clumps removed by gentile aggregation. The number of attached cells were either immediately determined or the cell coated stents were placed in a 60 mm dish containing complete culture media and cultured for the indicated time. To determine cell attachment, the stent was placed in 0.05% TE to dissociate the cell from the stent and the number of cells determined using standard hemocytometer technique. Complete cell removal from the stent was confirmed by microscopy. Cell numbers were normalized to precoated stent weights and represented as fold of unmodified control stents. Additionally, the percent flow diverter stent coverage was determined by measuring the open ie. non-cellular covered area within the intra-strut space divided by the total intra-strut area. A minimum of 6 microscopic images from various sides of the stent were obtained at 20x magnification and the open area measured using ImageJ. To determine the effects of surface coatings on cellular migration, endothelial cells were cultured in a 96 well dishes to 80-90% confluency and a flow diverter stent placed such that the end of the stent was in direct contact with the cellular layer. The culture plates were then returned to the cell culture incubator for the indicated times. Alternatively, flow diverter stents which were completely covered by endothelial cells were placed in a culture dish containing complete culture media but without cells and cellular migration from the stent to the plate determined.

### Rabbit Aneurysm Model

All animal procedures were approved by the Animal Care and Use Committee of the University of Miami. Aneurysms were created by intraluminal elastase treatment of the right common carotid artery (CCA) using digital subtraction angiography as previously described ^13,28,29^. Anesthesia was induced using 30 mg/kg Ketamine and 2 mg/kg Xylazine and maintained with isoflurane. To create the aneurysm, the right CCA was isolated and ligated distally. An arteriotomy was made proximally to the ligature, and a 4-Fr vascular sheath (Cordis, Miami Lakes, FL) was advanced to approximately 3 mm of the CCA origin. A 3-Fr Fogarty ballon (Baxter, Deerfield, IL) was then advanced through the sheath to the origin of the CCA and inflated to induce right CCA flow arrest. One milliliter of porcine elastase (approximately 130 U/ml, Worthington Biochemical) was incubated in the CCA lumen for 20 min. The elastase, balloon and sheath were then removed and the right CCA ligated below the sheath access site. Approximately four weeks following aneurysm creation, the right femoral artery was isolated, and a 6-Fr sheath placed. A 5-Fr Envoy guide catheter (Johnson and Johnson, Cincinnati, OH) was advanced to the right subclavian artery. A Phenom catheter was used to place two unmodified or modified flow diverter stent across the neck of the aneurysm. Aspirin (10 mg/kg) and clopidogrel (10 mg/kg) were given two days prior to device implantation and for the duration of the experiment. Aneurysm occlusion was determined 90 -100 days post-treatment by digital subtraction angiography of the aortic arch using femoral endovascular techniques and the Raymond-Roy Occlusion Score^30^. Raymond-Roy Occlusion scoring was conducted in a blinded fashion to stent coating. Animals were then euthanized with a lethal injection of euthazol and the device-bearing parent artery and aneurysm harvested and immediately placed in 10% neutral buffered formalin.

### Tissue Histology

The aneurysms and device-bearing parent arteries were fixed in 10% neutral buffered formalin for at least 24 hrs. The tissue was then processed for histological as previously described and cut into 4 μm sections^31^. Serial sections of each aneurysm and parent artery underwent HCE staining. Endothelial and vascular smooth muscle cell stent coverage across the aneurysm neck was determined by *en face* immunohistology analysis using primary antibodies against the endothelial cell maker, CD31 and the vascular smooth muscle cell marker, alpha-smooth muscle actin. Histological analysis of the HCE stained sections were performed by a blinded investigator for: (a) endothelial cell coverage; (b) inflammation; and (c) smooth muscle cell coverage^13^. The grading scale is shown in online supplementary data.

### Statistics

All data are expressed as mean ± S.E.M. Statistical analysis between two groups was carried out using the unpaired Student’s t-test. Statistical analysis between more than two groups was performed using a one-way ANOVA with Dunnett’s multiple comparison post hoc test. Statistical analysis of categorical variables was carried out using Chi-square and Fisher’s exact tests as appropriate. A p-value of less than 0.05 was considered statistically significant for all analyses.

## RESULTS

### Neuroform Stent Coating In Vivo

Flow diverting stents are primarily made of either nitinol, a nickel-titanium alloy or cobalt-chromium. Therefore, for our *in vitro* studies we tested the coating of nitinol -Neuroform stents and cobalt-chromium Pipeline flow diverters. Additionally, coating cell culture plates with fibronectin enhances endothelial attachment and overall health^32^. Therefore, for our *in vitro* studies we coated stents with fibronectin as a positive control. Initially we determined if coating Neuroform stents enhanced cell attachment. Therefore, stents were incubated with an endothelial cell suspension as detailed in the methods and the number of cells attaching to the stent determined immediately following cell-stent incubation. As shown in Figure 1A and B, endothelial cells were found attached to unmodified, and gelatin and fibronectin coated stents. Cells attached to control stents were rounded and not fully attached as indicated by a flat state. In contrast, on both gelatin and fibronectin coated stents the cells were primarily fully attached with only a few rounded cells observed. Interestingly, cells on fibronectin coated stents showed enhanced cell growth near points of strut narrowing. This was also observed in gelatin coated and uncoated stents but only after a few days in culture (Data not shown). Immediately following cell-stent incubation, the number of cells attached to gelatin coated stents was increased nearly 6-fold but was not significantly different from unmodified stents. In contrast, there was a 13-fold increase (p< 0.05) in cells attached to fibronectin coated stents, compared to unmodified controls. Endothelial cell coverage of the stents was extensive after 4 days in culture with a visible cellular layer covering the stent struts (Figure 1C). There was no difference in cell death as indicated by propidium iodide staining and no significant difference in cell numbers between gelatin and unmodified controls (Figure1 D and E).

**Figure 1.**
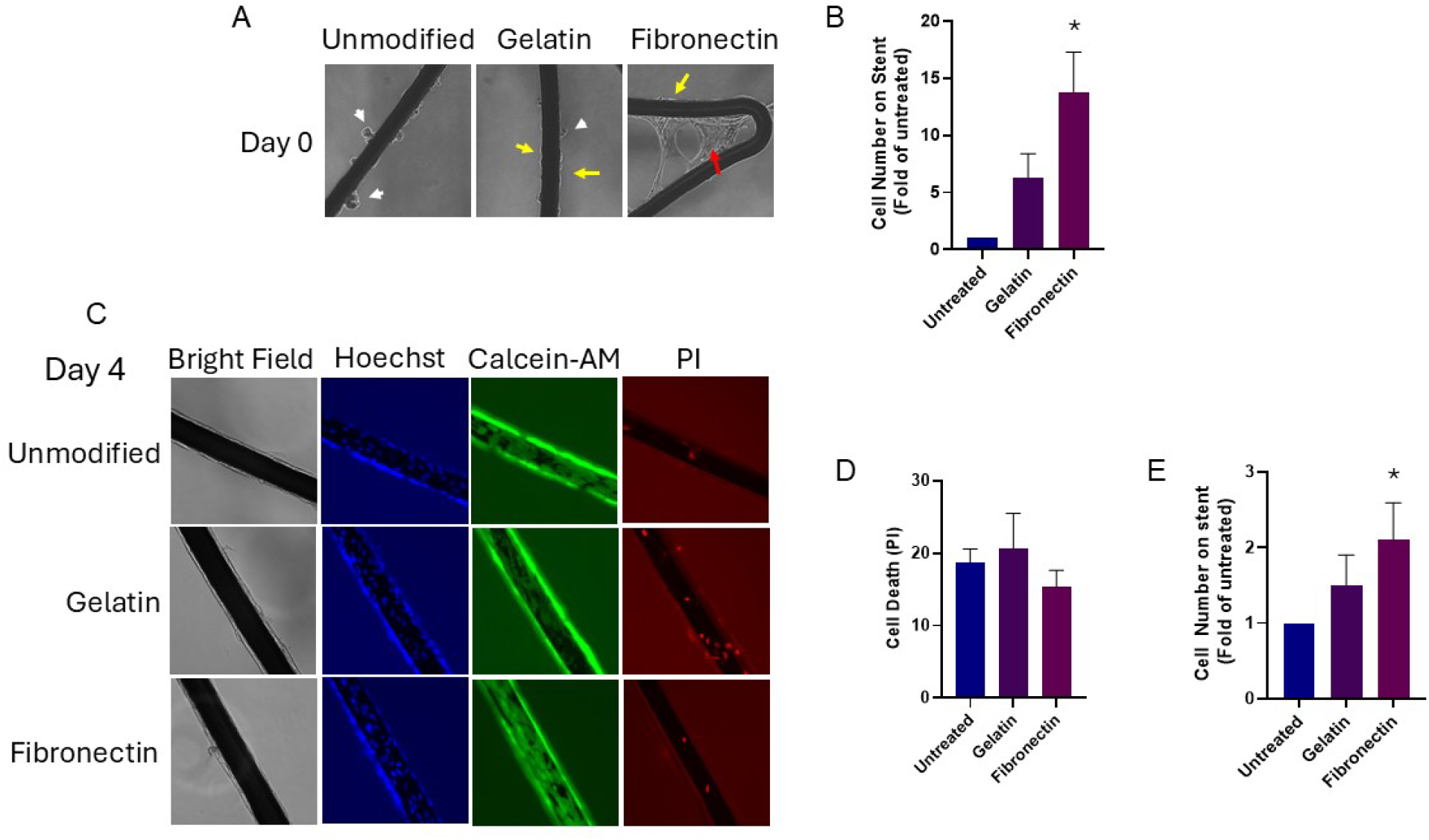
Fibrinogen but not gelatin coating increases cellular attachment to nitinol stents. (A) Representative micrographs of human brain endothelial cell attachment to Neuroform stents immediately following stent-cell incubation for 3 hours. White arrow heads indicate attached but rounded cells, yellow arrows indicate completely attached flattened cells, and red arrow indicates cellular growth and extension. Quantification of the number of cells attached to the stent is shown in (B, n = 4, * represents p < 0.05 versus unmodified control stents). In (C) representative bright-field, Hoechst, Calcein-AM, and propidium iodide (PI) micrograph showing cellular growth and stent coverage of the endothelial attached cells following 4 days of culture. Quantitation of PI positive cell death and cell numbers are shown in (D) and (E) respectively. n = 4, * represents p < 0.05 versus unmodified control stents.

### Flow Diverting Stent Coating In Vivo

Next, we determined cell attachment to gelatin and fibronectin coated flow diverter stents. Flow diverting stents are designed to act as a scaffold for cellular growth and coverage and as a result, only a few rounded cells were observed on unmodified or coated stents immediately following cell incubation (Figure 2A). Culturing the cell-seeded-stents for 5 days demonstrated significant (p< 0.05) cell proliferation on both gelatin (8.8 ± 2.8-fold) and fibronectin (15 ± 2.8-fold) coated stents compared to unmodified controls (Figure 2B). Further culturing of the endothelial cell seeded stent caused complete cellular coverage of the unmodified and modified stents with no evident cell death as indicated by propidium iodide staining (Figure 2C).

**Figure 2.**
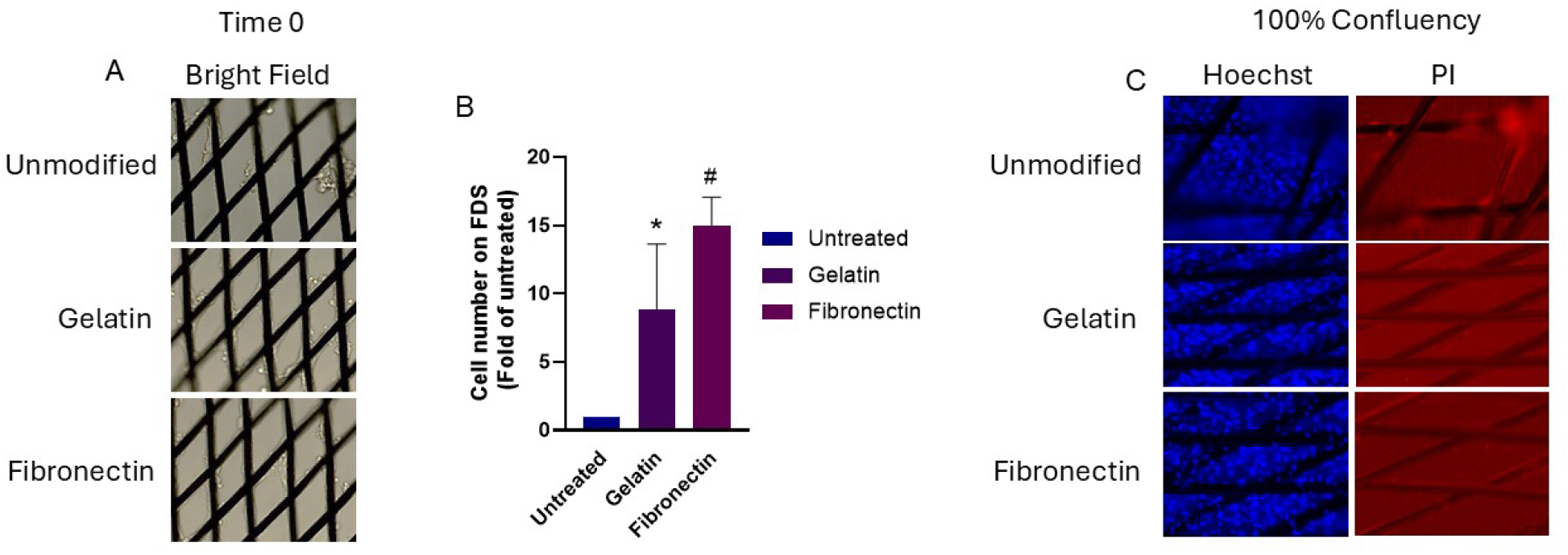
Gelatin coating of flow diverting stents enhances cellular attachment and proliferation. Representative brightfield micrographs of human brain endothelial cell attachment to unmodified and gelatin and fibronectin modified flow diverting stents immediately following stent-cell incubation for 3 hours (A). The number of cells on gelatin and fibronectin coated stents 5 days following cell seeding relative to unmodified controls is shown in (B). Representative micrograph of Hoechst and propidium iodide (PI) staining of fully reendothelized stents (C). n = 4, * represents p < 0.05 versus unmodified control stents.

While culturing the flow diverting stents, we observed differences in the rate of cellular coverage on the gelatin and fibronectin stents compared to controls. Therefore, we measured the rate of cellular coverage by determining changes in the non-cellularized area between the struts over a 22-day period. As shown in Figure 3A and B, 20% of gelatin coated stents and 11% of fibronectin stents were covered by cells after 3 days in culture. While only 3% of the unmodified stent was covered by cells. By 7 days in culture, nearly 75% of the gelatin coated stent (p< 0.001) and 53% of fibronectin stents were covered by endothelial cells. In contrast only 20% of the unmodified stent was covered. By two weeks, gelatin and fibronectin stents were 100% (p< 0.001) covered by endothelial cells, while unmodified stents were only 50% covered. It took an additional week before the unmodified stents were fully covered by endothelial cells.

**Figure 3.**
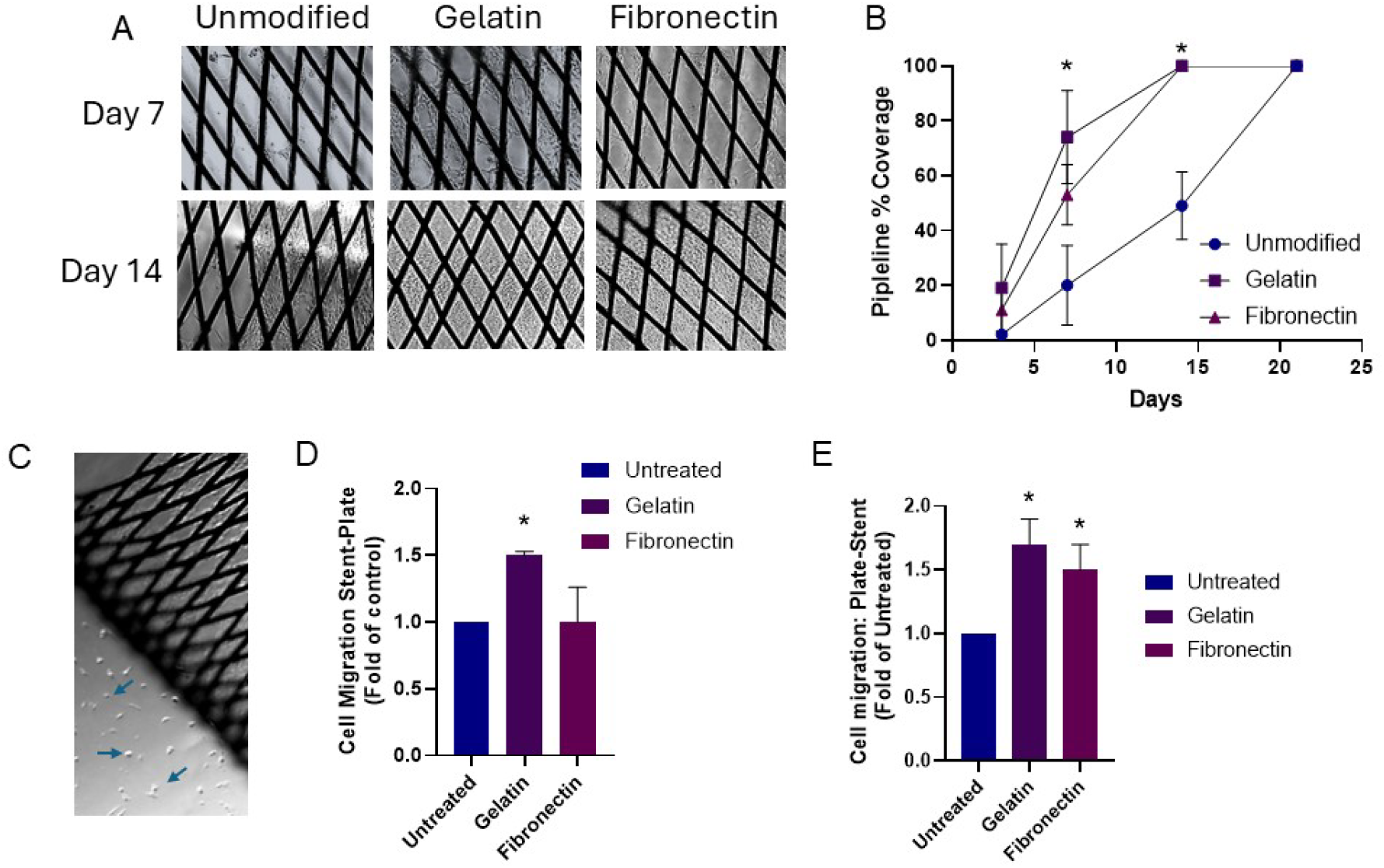
Gelatin coating enhances cellular coverage of flow diverting stents. Representative brightfield micrographs of human brain endothelial cell attachment to unmodified and gelatin and fibronectin modified flow diverting stents at 7 and 14 days following cell-stent incubation (A). Quantitation of endothelial stent coverage over 21 days is shown in (B). n= 5, * p < 0.001 versus unmodified control stents. In (C) representative bright filed microscopy image of cells which have migrated from a stent onto a cell culture dish. Quantitation of cell migration from the stent to the cell culture dish and from the culture dish to the stent is shown in (D and E), respectively. n= 3, * p < 0.05 versus unmodified control stents.

### Role of Gelatin Coating on Cell Migration

As we cultured the flow diverting stents over time, we also noticed the migration of cells from the stents to the cell culture plate. Therefore, we determined if gelatin and fibronectin coating increased cell migration. To test this, stents were coated with endothelial cells and cultured until reaching 100% confluency. The stents were then placed into a new culture dish without cells and the number of cells migrating from the stent to the plate determined after 7 days. As shown in Figure 3C, cellular migration from the stent to the plate was observed with each stent type. However, only gelatin modification significantly (p< 0.05) increased cell migration by nearly 50% compared to unmodified stents. To better mimic real-world conditions, coated and unmodified stents which were not seeded with cells were placed on a nearly confluent layer of endothelial cells and the migration of cells onto the stent determined after 7 days in culture. As shown in Figure 3E, cells were found to migrate onto modified as well as unmodified stents. However, gelatin stent modification significantly increased the number of cells migrating onto the stent by more than 50% (p< 0.05). Overall, these results demonstrate that gelatin coating only modestly enhances cell attachment to nitinol stents but significantly increases endothelial attachment and migration on cobalt-chromium alloy stents. Therefore, we were interested in determining if endothelial cell-seeded-gelatin-coated flow diverter stents improved aneurysm occlusion and parent vessel healing.

### Assessment of Gelatin Coating In Vivo

To test this, we created aneurysms in rabbits and allowed the aneurysm to mature for at least one month before treatment. Angiographic imaging prior to treatment demonstrated no difference in aneurysm height, width, and neck size or right brachiocephalic parent artery size (Table 1). The aneurysms were then treated with either unmodified or gelatin coated flow diverting stents seeded with GFP-expressing human aortic endothelial cells. Importantly, stent modification did not affect stent delivery, deployment or wall apposition. Representative angiographic images of pre-treatment, immediately post-treatment and pre-termination at 90-100 days is shown in Figure 4A. Angiographic assessment using the Raymond-Roy Occlusion Score found that aneurysms in unmodified cohorts were completed occluded in 4 of 7 animals (57%, RROC 1), while partial aneurysm filling was found in 2 of 7 animals (29%, RROC 2) and complete aneurysm filling found in 1of 7 animals (14%, RROC 3). In contrast, there was 100% aneurysm occlusion (8 of 8 animals, RROC 1, p< 0.001) treated with cell-seeded-gelatin modified stents (Figure 4B).

**Figure 4.**
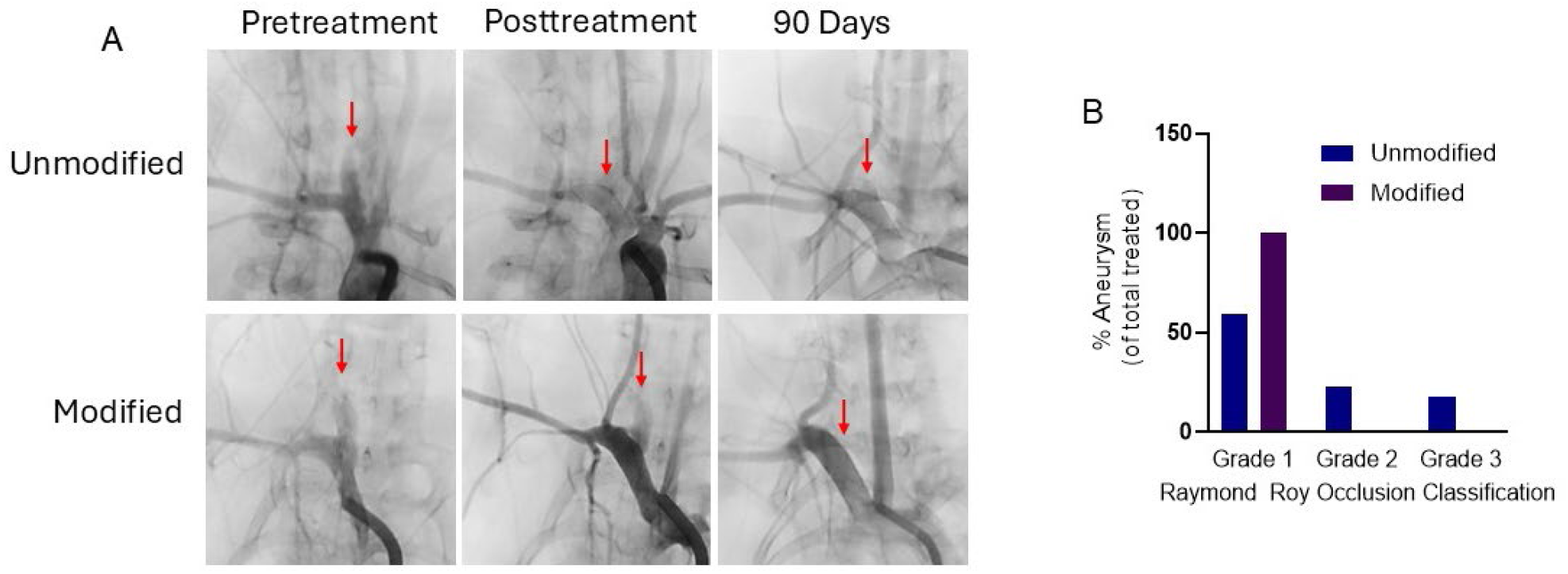
Stent modification enhances aneurysm occlusion. (A) Representative angiographic images showing aneurysm filling pre-implant, immediately post-implant, and prior to euthanasia at 90 days. (B) Raymond Roy Occlusion Classification at 90 days. Grade 1 designates complete occlusion, Grade 2 designates residual neck filling, and Grade 3 designates visible dome filling. p < 0.001 using Fisher’s exact test. N = 7 controls and 8 endothelial cell-seeded-gelatin coated modified stents.

**Table:** Average aneurysm dimensions and brachiocephalic trunk diameter measured prior to aneurysm treatment.

| Device | Aneurysms Neck (mm) | Aneurysm Height (mm) | Aneurysm Width (mm) | Right BCT (mm) |
| --- | --- | --- | --- | --- |
| Unmodified | 3.77 ± 0.77 | 7.63 ± 1.90 | 3.20 ± 0.43 | 3.27 ± 0.52 |
| Modified | 4.76 ± 2.17 | 8.05 ± 3.70 | 3.11 ± 0.41 | 3.23 ± 0.36 |

Histological assessment through the neck and aneurysm dome was assessed using HCE staining (Figure 5A-C), while luminal endothelial cells and vascular smooth muscle cell coverage across the aneurysm neck were assessed by immunohistochemistry (Figure 5D). Histological quantitation revealed no significant changes in overall endothelial and vascular smooth muscle cell neck coverage or inflammatory response (Figure 5E). However, there was a trend in increased inflammatory cells located around or near the struts of the modified stents. Quantitative analyses of neck neoarterial wall formation demonstrated a trend in increase wall thickness (16 ± 11 μm in unmodified and 34 ± 23 μm in modified stents). Similarly, neointimal formation in the right brachiocephalic artery revealed a nonsignificant increase in wall thickness (27 ± 13 μm in unmodified and 48 ± 30 μm in modified stents). However, angiographic images prior to sacrifice revealed strong perfusion of the parent artery as well as side branch arteries such as the right vertebral artery in modified stent treated cohorts (Figure 6A and B). Immunocytochemistry of modified stents reveals strong endothelial cell stent coverage but not in the region of the side branching artery

**Figure 5.**
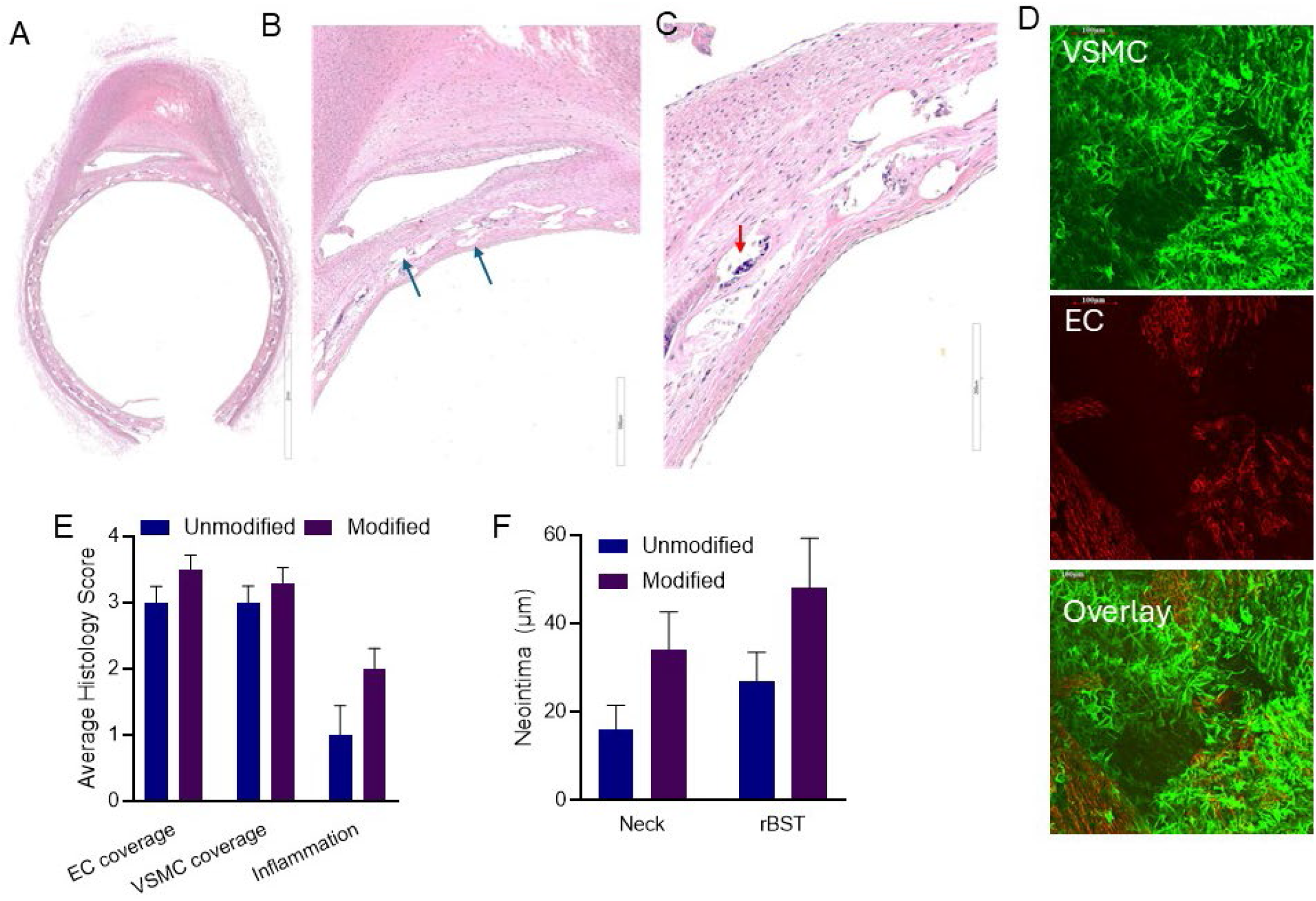
Representative histology of neoarterial wall formation 90 days following aneurysm treatment. In (A), HCE staining of a healed aneurysm treated with gelatin coated stent. Blue arrows represent examples of strut location after removal (B). Red arrow represents an example of inflammatory cells in the location of the stent (C). In (D), endothelial (EC) and vascular smooth muscle (VSMC) immunohistochemistry staining of the intraluminal surface of the neo-arterial neck wall. Average histological assessment of EC and VSMC coverage and inflammation response is shown in (E). Quantitative analysis of neo-arterial (Neck) and right brachiocephalic trunk (rBST) neointimal formation is shown in (F).

**Figure 6.**
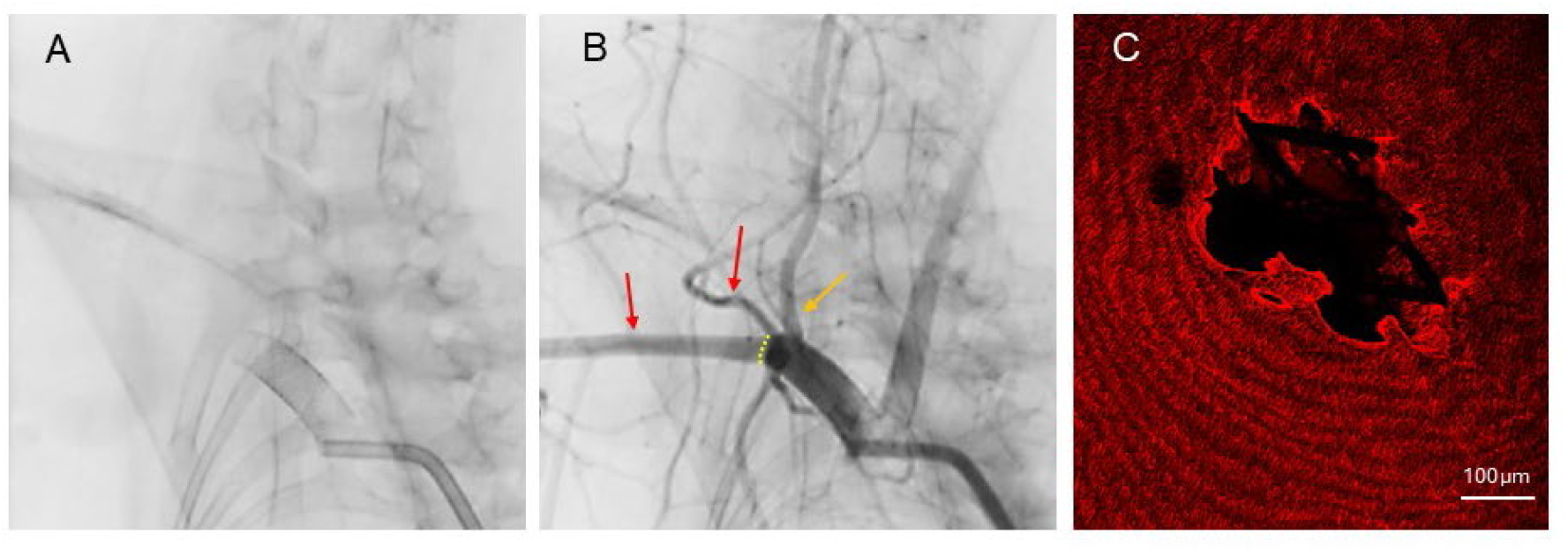
Parent vessel perfusion and side branch patency are not altered by gelatin coating. Representative angiographic image of a gelatin coated stent location at 90 days following treatment(A). DSA image of parent and side branch patency is shown in (B). Yellow dotted line represents distal location of the stent end. Red arrows demonstrate subclavian and branching artery perfusion. Yellow arrow represents right vertebral artery covered by the deployed stent. In (C) intraluminal surface endothelial cell staining demonstrates patent vertebral artery shown in (B)

## DISCUSSION

In this study we demonstrate the therapeutic potential of gelatin as a stent surface coating to enhance endothelial cell coverage and increase aneurysm obliteration and parent vessel healing. Our results demonstrate that 1) gelatin coating enhances endothelial cell attachment, proliferation, migration and stent coverage which was dependent upon the metal composition of the stent and 2) endothelial cell seeded-gelatin coated-flow diverting stents caused complete aneurysm obliteration and neoarterial formation without affecting side branch patency or parent artery perfusion.

With the advancement of endovascular treatment options, the clinical outcomes of cerebral aneurysm patients have improved over the past decade^33,34^. However, aneurysm recanalization, thrombotic complications, and delayed rupture remain important causes of morbidity^5,35,36^. Following flow diversion stent placement, neointimal and neoartrial wall formation across the aneurysm neck is associated with thrombotic complications that can lead to stroke and allows continued aneurysm filling^5,37^. Therefore, increasing the rate of neointimal development would accelerate parent vessel healing and thereby exclude the aneurysm and stent from the circulation and prevent thrombotic complications and possible aneurysm rupture. As a result, research has primarily focused on stent coatings to reduce thrombogenicity while enhancing cellularization of the stent^8,9,38^.

Gelatin is derived from the partial hydrolysis of collagen, which constitutes the main structure of the extracellular matrix^39,40^. Gelatin is one of the most widely used natural biomaterials in medicine and has excellent biocompatibility, biodegradability, and low immunogenicity^39^. It contains bioactive motifs like the RGD (arginyl-glycyl-aspartic acid) sequence that promote cell adhesion, spreading, and proliferation^39–42^. Although used in medical applications, there has been limited use of gelatin in neurovascular applications, particularly flow diverting stent coatings.

Flow diverting stents are primarily made of nitinol a nickel-titanium alloy and cobalt-chromium alloy, with radiopaque elements such as platinum-tungsten wires or tantalum interwoven for enhanced fluoroscopic visualization^1^. Our results demonstrate that the stent metal composition affects the potential of the gelatin coating. On nitinol stents gelatin coating was found to enhance the rate of endothelial cell attachment as indicated by flattened versus rounded cell morphology following cell-stent incubation, but it did not affect overall cell attachment numbers, proliferation or stent coverage. In contrast gelatin coating of cobalt-chromium stents greatly increased endothelial cell attachment, proliferation and migration. Interestingly, we found that gelatin coating also increased the rate of stent endothelial cell coverage by 33% such that gelatin coated stents were completely endothelized by 2 weeks while unmodified stents were only completely covered by 3 weeks. In contrast to our findings, Meer et al.^43^ found that gelatin coating of nitinol stents could enhance endothelial cell attachment when incubated with cells for 16-24 hours, which is a much longer incubation period than the 3 hours used in this study. Similarly, Peng et al.^44^ found increased mesenchymal stem cell attachment to UV treated-nitinol stents coated with 8% gelatin/ polylysine mixture and incubated with cells for 24 hours or more.

An alternative approach to enhancing parent vessel healing and neointima formation is the delivery or recruitment of cells to the stent. For example, coating stents with CD34 and CD133 antibodies to capture circulating endothelial progenitor cells improved cell attachment and stent coverage by endothelial progenitor cells both in culture and when implanted the rabbit descending aorta^16,45^. Likewise, coating flow diverting stents with a CD31-mimetic peptide increased endothelial cell attachment and coverage and enhanced aneurysm occlusion in a rabbit aneurysm model^21^. However, in this study we were interested in taking a different approach and determining if stents seeded with endothelial cells would enhance aneurysm obliteration and neointima formation. Although the endothelial cells were stably transfected with GFP expression we were unable to identify any GFP cells in the treated vessel. However importantly, we found that endothelial cell - seeded-gelatin-coated-flow diverter stents resulted in 100% aneurysm obliteration which is a 41% increase in obliteration rate compared to controls at 90 days. Immunohistochemistry analysis demonstrated complete endothelial cell and vascular smooth muscle cell neck coverage. Histologically, there were no differences in endothelial cell and vascular smooth muscle cell coverage, inflammation or neointimal formation compared to controls. However, there were trends to increased inflammatory cells located around the stent’s struts, and increased neointima formation at the aneurysm neck and parent artery. Importantly, cell-seeding did not affect parent and distal vessel perfusion or side branch patency. Although there are studies investigating the potential of gelatin coating for endothelial progenitor cells and mesenchymal stem cell stent seeding^16,44–46^, we believe this is the first study in which cells were seeded onto a flow diverting stent for aneurysm treatment.

There are several limitations to this study. We investigated aneurysm occlusion and vessel healing at approximately 90 days posttreatment, so long-term durability as well as early beneficial effects of endothelial cell-seeded and gelatin coating on parent vessel healing were not investigated. Therefore, aneurysm occlusion rates at both early and late time points should be investigated. Additionally, to monitor the transplanted cells *in vivo* stably expressing GFP endothelial cells were seeded on gelatin coated stents. However, GFP positive cells were not identified in the harvested tissues. Therefore, it is unclear how long the seeded cells survived after transplantation and what effect they had on enhancing aneurysm occlusion vessel healing. Further time points and gelatin coating alone without seeded cells could help define this in future studies.

## CONCLUSIONS

We demonstrate that gelatin coating of cobalt-chromium stents improves endothelial cell attachment and the rate of cellular coverage. Consistent with this, we found that endothelial seeded gelatin stents resulted in complete aneurysm occlusion and endothelialization across the aneurysm neck. The gelatin coating procedure used in this study is suitable for scaling-up and potential use in patients. Thus, gelatin coating may be a promising strategy to increase stent cellularization and parent vessel healing which could improve aneurysm treatment.

## REFERENCES

1. Becske T, Kallmes DF, Saatci I, et al. Pipeline for uncoilable or failed aneurysms: results from a multicenter clinical trial. Radiology 2013;267:858–68.

2. Hanel RA, Kallmes DF, Lopes DK, et al. Prospective study on embolization of intracranial aneurysms with the pipeline device: the PREMIER study 1 year results. J Neurointerv Surg 2020;12:62–6.

3. Brinjikji W, Murad MH, Lanzino G, Cloft HJ, Kallmes DF. Endovascular treatment of intracranial aneurysms with flow diverters: a meta-analysis. Stroke 2013;44:442–7.

4. Becske T, Brinjikji W, Potts MB, et al. Long-Term Clinical and Angiographic Outcomes Following Pipeline Embolization Device Treatment of Complex Internal Carotid Artery Aneurysms: Five-Year Results of the Pipeline for Uncoilable or Failed Aneurysms Trial. Neurosurgery 2017;80:40–8.

5. Kallmes DF, Hanel R, Lopes D, et al. International retrospective study of the pipeline embolization device: a multicenter aneurysm treatment study. AJNR Am J Neuroradiol 2015;36:108–15.

6. Saber H, Kherallah RY, Hadied MO, Kazemlou S, Chamiraju P, Narayanan S. Antiplatelet therapy and the risk of ischemic and hemorrhagic complications associated with Pipeline embolization of cerebral aneurysms: a systematic review and pooled analysis. J Neurointerv Surg 2019;11:362–6.

7. Brinjikji W, Lanzino G, Cloft HJ, Siddiqui AH, Kallmes DF. Risk Factors for Hemorrhagic Complications following Pipeline Embolization Device Treatment of Intracranial Aneurysms: Results from the International Retrospective Study of the Pipeline Embolization Device. AJNR Am J Neuroradiol 2015;36:2308–13.

8. Hagen MW, Girdhar G, Wainwright J, Hinds MT. Thrombogenicity of flow diverters in an ex vivo shunt model: effect of phosphorylcholine surface modification. J Neurointerv Surg 2017;9:1006–11.

9. Girdhar G, Andersen A, Pangerl E, et al. Thrombogenicity assessment of Pipeline Flex, Pipeline Shield, and FRED flow diverters in an in vitro human blood physiological flow loop model. J Biomed Mater Res A 2018;106:3195–202.

10. Zoppo CT, Mocco J, Manning NW, Bogdanov AA, Jr., Gounis MJ. Surface modification of neurovascular stents: from bench to patient. J Neurointerv Surg 2024;16:908–13.

11. King RM, Peker A, Epshtein M, et al. Active drug-coated flow diverter in a preclinical model of intracranial stenting. J Neurointerv Surg 2024;16:731–6.

12. Bricout N, Chai F, Sobocinski J, et al. Immobilisation of an anti-platelet adhesion and anti-thrombotic drug (EP224283) on polydopamine coated vascular stent promoting anti-thrombogenic properties. Mater Sci Eng C Mater Biol Appl 2020;113:110967.

13. Starke RM, Thompson J, Pagani A, et al. Preclinical safety and efficacy evaluation of the Pipeline Vantage Embolization Device with Shield Technology. J Neurointerv Surg 2020;12:981–6.

14. Zoppo CT, Epshtein M, Gounis MJ, Anagnostakou V, King RM. Longitudinal healing flow diverting stents with phosphorylcholine surface modification. J Neurointerv Surg 2024;16:582–6.

15. Hackett AM, Luther EM, Walker AP, et al. Telescoping pipeline vantage embolization devices with shield technology for the treatment of a giant, symptomatic dolichoectatic basilar trunk aneurysm. Surg Neurol Int 2022;13:434.

16. Aoki J, Serruys PW, van Beusekom H, et al. Endothelial progenitor cell capture by stents coated with antibody against CD34: the HEALING-FIM (Healthy Endothelial Accelerated Lining Inhibits Neointimal Growth-First In Man) Registry. J Am Coll Cardiol 2005;45:1574–9.

17. Duckers HJ, Soullié T, den Heijer P, et al. Accelerated vascular repair following percutaneous coronary intervention by capture of endothelial progenitor cells promotes regression of neointimal growth at long term follow-up: final results of the Healing II trial using an endothelial progenitor cell capturing stent (Genous R stent). EuroIntervention 2007;3:350–8.

18. Marosfoi M, Langan ET, Strittmatter L, et al. In situ tissue engineering: endothelial growth patterns as a function of flow diverter design. J Neurointerv Surg 2017;9:994–8.

19. Diaz-Rodriguez S, Rasser C, Mesnier J, et al. Coronary stent CD31-mimetic coating favours endothelialization and reduces local inflammation and neointimal development in vivo. Eur Heart J 2021;42:1760–9.

20. Woodfin A, Voisin MB, Nourshargh S. PECAM-1: a multi-functional molecule in inflammation and vascular biology. Arterioscler Thromb Vasc Biol 2007;27:2514–23.

21. Cortese J, Rasser C, Even G, et al. CD31 Mimetic Coating Enhances Flow Diverting Stent Integration into the Arterial Wall Promoting Aneurysm Healing. Stroke 2021;52:677–86.

22. Echave MC, Saenz del Burgo L, Pedraz JL, Orive G. Gelatin as Biomaterial for Tissue Engineering. Curr Pharm Des 2017;23:3567–84.

23. Rizwan M, Yao Y, Gorbet MB, et al. One-Pot Covalent Grafting of Gelatin on Poly(Vinyl Alcohol) Hydrogel to Enhance Endothelialization and Hemocompatibility for Synthetic Vascular Graft Applications. ACS Appl Bio Mater 2020;3:693–703.

24. Klotz BJ, Gawlitta D, Rosenberg A, Malda J, Melchels FPW. Gelatin-Methacryloyl Hydrogels: Towards Biofabrication-Based Tissue Repair. Trends Biotechnol 2016;34:394–407.

25. Su K, Wang C. Recent advances in the use of gelatin in biomedical research. Biotechnol Lett 2015;37:2139–45.

26. Xing Y, Gu Y, Guo L, et al. Gelatin coating promotes in situ endothelialization of electrospun polycaprolactone vascular grafts. J Biomater Sci Polym Ed 2021;32:1161–81.

27. Kopeć K, Wojasiński M, Eichler M, et al. Polydopamine and gelatin coating for rapid endothelialization of vascular scaffolds. Biomater Adv 2022;134:112544.

28. Dai D, Bilgin C, Ding Y, et al. How the elastase-induced rabbit aneurysm heals following flow diverter treatment: a histopathological study. J Neurosurg 2024;141:1262–9.

29. Thompson JW, Elwardany O, McCarthy DJ, et al. In vivo cerebral aneurysm models. Neurosurg Focus 2019;47:E20.

30. Roy D, Milot G, Raymond J. Endovascular treatment of unruptured aneurysms. Stroke 2001;32:1998–2004.

31. Dai D, Ding YH, Danielson MA, et al. Modified histologic technique for processing metallic coil-bearing tissue. AJNR Am J Neuroradiol 2005;26:1932–6.

32. Mancianti ML, Fimiani M, Castelli A, Raffaelli M, Perotti R, Valentino A. [Endothelial cell culture as a model for the study of wound healing]. Boll Soc Ital Biol Sper 1984;60:473–8.

33. Eskey CJ, Meyers PM, Nguyen TN, et al. Indications for the Performance of Intracranial Endovascular Neurointerventional Procedures: A Scientific Statement From the American Heart Association. Circulation 2018;137:e661–e89.

34. Molyneux AJ, Kerr RS, Yu LM, et al. International subarachnoid aneurysm trial (ISAT) of neurosurgical clipping versus endovascular coiling in 2143 patients with ruptured intracranial aneurysms: a randomised comparison of effects on survival, dependency, seizures, rebleeding, subgroups, and aneurysm occlusion. Lancet 2005;366:809–17.

35. Ferns SP, Sprengers ME, van Rooij WJ, et al. Late reopening of adequately coiled intracranial aneurysms: frequency and risk factors in 400 patients with 440 aneurysms. Stroke 2011;42:1331–7.

36. Kulcsár Z, Houdart E, Bonafé A, et al. Intra-aneurysmal thrombosis as a possible cause of delayed aneurysm rupture after flow-diversion treatment. AJNR Am J Neuroradiol 2011;32:20–5.

37. Li ZF, Fang XG, Yang PF, et al. Endothelial progenitor cells contribute to neointima formation in rabbit elastase-induced aneurysm after flow diverter treatment. CNS Neurosci Ther 2013;19:352–7.

38. Hellstern V, Aguilar Pérez M, Henkes E, et al. Use of a p64 MW Flow Diverter with Hydrophilic Polymer Coating (HPC) and Prasugrel Single Antiplatelet Therapy for the Treatment of Unruptured Anterior Circulation Aneurysms: Safety Data and Short-term Occlusion Rates. Cardiovasc Intervent Radiol 2022;45:1364–74.

39. Young S, Wong M, Tabata Y, Mikos AG. Gelatin as a delivery vehicle for the controlled release of bioactive molecules. J Control Release 2005;109:256–74.

40. Ahmady A, Abu Samah NH. A review: Gelatine as a bioadhesive material for medical and pharmaceutical applications. Int J Pharm 2021;608:121037.

41. Ruoslahti E, Pierschbacher MD. Arg-Gly-Asp: a versatile cell recognition signal. Cell 1986;44:517–8.

42. Hersel U, Dahmen C, Kessler H. RGD modified polymers: biomaterials for stimulated cell adhesion and beyond. Biomaterials 2003;24:4385–415.

43. Ter Meer M, Daamen WF, Hoogeveen YL, et al. Continuously Grooved Stent Struts for Enhanced Endothelial Cell Seeding. Cardiovasc Intervent Radiol 2017;40:1237–45.

44. Peng Q, Guo R, Zhou Y, Teng R, Cao Y, Mu S. Comparison of Gelatin/Polylysine- and Silk Fibroin/SDF-1α-Coated Mesenchymal Stem Cell-Seeded Intracranial Stents. Macromol Biosci 2023;23:e2200402.

45. Wu X, Yin T, Tian J, et al. Distinctive effects of CD34- and CD133-specific antibody-coated stents on re-endothelialization and in-stent restenosis at the early phase of vascular injury. Regen Biomater 2015;2:87–96.

46. Larsen K, Cheng C, Tempel D, et al. Capture of circulatory endothelial progenitor cells and accelerated re-endothelialization of a bio-engineered stent in human ex vivo shunt and rabbit denudation model. Eur Heart J 2012;33:120–8.

